# Obg-like ATPase 1 Genetic Deletion Leads to Dilated Cardiomyopathy in Mice and Structural Changes in Drosophila Heart

**DOI:** 10.1101/2024.05.28.596265

**Authors:** Praveen K Dubey, Sarojini Singh, Hussain Khalil, Goutham K Kommini, Krishna Moorthi Bhat, Prasanna Krishnamurthy

**Affiliations:** Department of Biomedical Engineering, Schools of Medicine and Engineering, University of Alabama at Birmingham, AL, USA; Department of Molecular Medicine, University of South Florida, Tampa, Florida, USA

**Keywords:** OLA1, dilated cardiomyopathy, heart failure, LV dysfunction

## Abstract

Cardiomyopathy, disease of the heart muscle, is a significant contributor to heart failure. The pathogenesis of cardiomyopathy is multifactorial and involves genetic, environmental, and lifestyle factors. Identifying and characterizing novel genes that contribute to cardiac pathophysiology are crucial for understanding cardiomyopathy and effective therapies.

In this study, we investigated the role of a novel gene, *Obg-like ATPase 1* (*Ola1*), in cardiac pathophysiology using a cardiac-specific knockout mouse model as well as a Drosophila model. Our previous work demonstrated that OLA1 modulates the hypertrophic response of cardiomyocytes through the GSK-beta/beta-catenin signaling pathway. Furthermore, recent studies have suggested that OLA1 plays a critical role in organismal growth and development. For example, *Ola1* null mice exhibit increased heart size and growth retardation. It is not known, however, if loss of function for *Ola1* leads to dilated cardiomyopathy.

We generated cardiac-specific *Ola1* knockout mice (OLA1-cKO) to evaluate the role of OLA1 in cardiac pathophysiology. We found that *Ola1*-cKO in mice leads to dilated cardiomyopathy (DCM) and left ventricular (LV) dysfunction. These mice developed severe LV dilatation, thinning of the LV wall, reduced LV function, and, in some cases, ventricular wall rupture and death. In Drosophila, RNAi-mediated knock-down specifically in developing heart cells led to the change in the structure of pericardial cells from round to elongated, and abnormal heart function. This also caused significant growth reduction and pupal lethality. Thus, our findings suggest that OLA1 is critical for cardiac homeostasis and that its deficiency leads to dilated cardiomyopathy and dysfunction. Furthermore, our study highlights the potential of the *Ola1* gene as a therapeutic target for dilated cardiomyopathy and heart failure.

## Introduction

Cardiomyopathy is a heterogeneous group of diseases that affect the structure and function of the heart muscle, resulting in significant morbidity and mortality worldwide (1). Dilated cardiomyopathy (DCM) is the most common type of cardiomyopathy and is characterized by enlargement of the heart chambers, thinning of the heart muscle, and reduced pumping function. This can lead to shortness of breath, fatigue, and swelling in the legs and ankles. DCM is a significant contributor to heart failure, accounting for up to one-third of cases of heart failure in developed countries. DCM is more common in men than in women, and the incidence increases with age (2,3).

The pathogenesis of DCM is multifactorial and involves genetic, environmental, and lifestyle factors. Inherited forms of DCM are caused by mutations in genes encoding proteins involved in the structural integrity and function of the heart muscle, such as sarcomeric proteins, cytoskeletal proteins, and ion channels. However, the majority of DCM cases are idiopathic or acquired, with no clear genetic cause (3).

Identifying and characterizing new genes that contribute to cardiac pathophysiology are crucial for understanding the development of DCM and designing effective therapeutic strategies. Several studies have reported genetic variants associated with DCM, including *titin*, *MYH7*, and *LMNA etc* (4). However, the genetic etiology of DCM remains poorly understood.

OBG-like ATPase 1 (OLA1) is a translation-factor-related (TRAFAC) class, YchF subfamily of P-loop GTPases, and *obg* family gene. It possesses both GTPase and ATPase activities and is involved in the translational regulation of cell proliferation, growth, and protein synthesis in mammals and yeast (5–7). Studies have shown that OLA1 is critical for normal mammalian development and that mice null for *ola1* exhibit growth retardation and developmental delay due to reduced cell proliferation rates (5). Our previous in vitro work demonstrated that OLA1 modulates the hypertrophic response of cardiomyocytes through the GSK-beta/beta-catenin signaling pathway (8). Interestingly, we also found a genetic mutation in the *Ola1* gene that is associated with human heart failure (9). However, its role in cardiac pathophysiology remains unknown.

The significance of this study lies in the potential role of OLA1 as a therapeutic target for DCM and heart failure. Currently, there are limited treatment options for DCM, and heart transplantation is often the only option for end-stage disease. Thus, the identification of novel therapeutic targets is essential to improve outcomes for patients with DCM and heart failure.

In this study, we aimed to investigate the role of a novel gene, *Ola1*, in cardiac pathophysiology using a cardiac-specific knockout mouse model. We observed that our cardiac-specific *Ola1* knockout mice spontaneously developed severe LV dilatation, thinning of the LV wall, reduced LV function, and, in some cases, ventricular wall rupture and death. These findings indicate that OLA1 is critical for cardiac homeostasis and that its deficiency leads to DCM and dysfunction. Additionally, we also used the Drosophila model system to investigate the effects of cardiac cell-specific loss of function of the Drosophila *ola1* gene on heart development. We found that the results from the fly system/gene were consistent with the findings in mice, with flies exhibiting both growth and cardiac issues.

In conclusion, our study suggests that OLA1 is critical for cardiac homeostasis, and its deficiency leads to DCM and dysfunction. It also highlights the role of *ola1* in cardiac development and function. Studies in our laboratory is on-going to elucidate the precise molecular mechanisms underlying the role of OLA1 in cardiac pathophysiology and DCM.

## Materials and methods

### Generation of cardiomyocyte specific OLA1 Knockout mice and genotyping

All animal procedures were approved by the Institutional Animal Care and Use Committee (IACUC) of the University of Alabama at Birmingham. To generate a cardiac-specific OLA1 knockout mice, we bred OLA1^fl/fl^ (B6(SJL)-Ola1^tm1.1Zshi^/J (stock no. 029493, Jackson Lab, USA) with α-myosin-Cre (B6.FVB-Tg(Myh6-cre)2182Mds/J; stock no. 011038, Jackson Lab, USA) from Jackson laboratories, USA. Genotyping was performed using Jackson laboratory protocols with minor modifications. DNA was isolated from ears punches. Ten microliters of amplified PCR products were run on a 1.5% agarose gel, and images were captured using a Bio-Rad Gel doc with a UV transilluminator.

For the survival study, animals were kept with standard food and water in a standard vivarium with a 12-hour light/12-hour dark cycle. Animals were observed for 80 weeks, and mortality was recorded during the observation period.

### Echocardiography

Transthoracic echocardiography on mice was performed using high-resolution micro-imaging systems (Vevo2100, Visual Sonics, Canada), equipped with a linear array transducer for mouse heart image acquisition. Briefly, mice were anesthetized (1–1.5% isoflurane) and allowed to breathe spontaneously (98.5–99% O_2_). They were placed supine on a temperature-controlled heating platform to maintain their body temperature at approximately 37°C. Parasternal short-axis M-mode tracings of the left ventricle (LV) were recorded to measure LV internal diameter in end-diastole (LVIDd) and calculate the LV ejection fraction (EF). Parasternal long-axis B-mode tracings of the right ventricular outflow tract (RVOT) were recorded to measure the RV area.

### Histological analysis

At the endpoint, animals were euthanized, and the hearts were immediately removed and fixed in 10% neutral-buffered formalin. Heart sections of 5 μm thickness were prepared, stained with hematoxylin and eosin and Masson’s trichrome, and imaged under a microscope (Nikon Eclipse E200 microscope) using NIS-Elements software version 4.60.

### Immunofluorescence imaging heart section

Paraffin-embedded tissue sections were deparaffined and permeabilized with PBS containing 0.1% Triton-X100 (PBST) at room temperature for 15 minutes. The sections were washed three times with PBS and blocked with PBST containing 2% goat serum albumin for 1 hour at room temperature. Next, the sections were incubated overnight at 4°C with primary antibodies against OLA1 (Proteintech), Cardiac Troponin T (Thermofisher, MA5-12960) at a 1:200 dilution in BSA. The sections were washed four times with PBS and then incubated for 1 hour with secondary antibodies. After four washes with PBS, the nuclei were stained with DAPI, and the sections were fixed in Vector Shield mounting media. All images were acquired using a fluorescent microscope.

### Fly stocks and Genetics

The following fly stocks were used: *tinc*GFP;*tinc*GAL4 and CG1354 RNAi (ola1RNAi). All crosses were done and maintained at 25°C. Canton S wild-type flies were used as control. Standard genetics were used.

### Pupal Lethality Assay

*tinc*GFP;*tinc*GAL4 virgins were separately crossed with Canton S and CG1354 RNAi males in 25°C. Normal and dead pupae were counted in both tinC>>ola1 RNAi cross and the control. Percentages of dead pupae were calculated. The experiment was repeated several times.

### Larval Heart Dissection and Imaging

Larvae were dissected within an artificial hemolymph solution (AHL). Four separate stock cocktail solutions (A-D) were made and stored at 4° for up to 60 days. These stock solutions were combined to prepare working AHL as follows: 10 mL of Solution A (1.08 M NaCl, 0.08 M MgCl_2_, 0.05 M KCl, 0.02 M 2H_2_O.CaCl_2_, 0.01 M NaH_2_PO_4_, 0.05 M HEPES), 10 mL of Solution B (0.1 M Sucrose), 5 mL of Solution C (0.1 M Trehalose dihydrate), and 1.6 mL of Solution D (0.25 M NaHCO_3_). Solution A, C, and D were autoclave sterilized and Solution B was filter sterilized. Upon preparation, AHL was pH adjusted to 7.1 and aerated for 30 minutes using an air stone and pump. The procedure to dissect and record live heart was adopted from Cooper at al. (2009). Briefly, larvae were placed on a Sylgard Petri dish, ventral side up to obscure tracheal tubes. Larvae were stabilized at both ends using insect pins placed between the posterior and anterior spiracles, with gentle stretching to ensure the body was extended. Approximately 200 µL of AHS. was pipetted onto the larva to prevent desiccation and to facilitate dissection. A small incision was made at the posterior end using a dissection needle to allow egress of internal organs. The body of the larva was slit open from the posterior to the anterior end using curved micro scissors, taking care to cut only the cuticle and avoid deeper structures. Exposed organs and fat bodies were carefully removed to clear the view of the heart tube using forceps and additional pins to manipulate the body flaps. Remaining connections between the heart tube and any organs were severed carefully to prevent displacement or damage to the heart. The preparation was rinsed repeatedly with fresh AHS to remove debris and waste products. The dissected heart was allowed to rest to ensure clear visibility and stabilization of the heart structures before observation and recording under an ApoTome microscope.

### RNA isolation and Quantitative Real-Time PCR

Total RNA was extracted from mice heart tissues using the TRIzol method (Invitrogen), followed by purification with the Qiagen RNA extraction kit (cat# 74106, Qiagen), in accordance with the manufacturer’s instructions. RNA quality and quantity were measured using a Nanodrop spectrophotometer. An equal amount (1ug) of RNA from each sample was reverse-transcribed using the RevertAid First Strand cDNA Synthesis Kit (cat# K1691, ThermoFisher Scientific). qPCR reactions were performed in a QuantStudio 3 system (Applied Biosystems, ThermoFisher Scientific) using PowerUp™ SYBR™ Green Master Mix (cat# A25778, ThermoFisher Scientific) and gene-specific primers. Target gene expression was normalized to housekeeping gene 18S rRNA and presented as fold change relative to the control.

### Western Blotting

Tissue lysates were prepared in RIPA buffer supplemented with a Thermofisher protease inhibitor cocktail. Protein concentration was determined using the Pierce BCA Protein Assay (Cat# 23235, Thermofisher), following the manufacturer’s protocol. Equal amounts of protein from each sample were denatured at 95°C in 4x Laemmli buffer, run on denaturing SDS-PAGE gels (4%-20%, Mini-PROTEAN TGX™ stain-free precast gels, Bio-Rad), and transferred to a PVDF membrane (Trans-Blot Turbo™ PVDF transfer system, Bio-Rad) using Bio-Rad’s standard protocol. The membrane was blocked with 5% non-fat dry milk in TBS-T and incubated overnight with primary antibodies OLA1 and GAPDH. This was followed by incubation with horseradish peroxidase (HRP)-conjugated secondary antibodies against rabbit and mouse. Visualization of the blots was achieved using the enhanced chemiluminescence (Pierce) detection system.

### Statistical analyses

All data are presented as mean ± standard error of the mean (SEM). An unpaired *t*-test was performed to determine statistical significance between the two groups using GraphPad Prism software version 10.2.0 (GraphPad Software Inc). A p-value of <0.05 was considered statistically significant.

## Results

### Generation and confirmation of Cardiac-specific *Ola1* knockout mice (*Ola1*-cKO)

OLA1 plays an essential role in various cellular processes and homozygous deletions of ola1 causes embryonic lethality, with a few escapers that are small in size and have enlarged hearts (5,10). In our recent study, we found that myocardial expression of OLA1 levels is significantly downregulated in human failing hearts (9). Our previous in vitro study demonstrated that OLA1 modulates the response to ANG II treatment in cardiomyocytes via the GSK3β/β-Catenin pathway (8). The flox mice were originally created by adding loxP sites flanking exon 2 of the *Ola1* gene, along with Frt sites flanking the neomycin resistance (Neo) cassette (5) (also shown in Figure 1A). To investigate the role of OLA1 in cardiac pathophysiology in vivo, we generated cardiomyocyte-specific *Ola1* knockout (cKO) mice by crossing *Ola1* flox/flox mice with transgenic mice expressing Cre recombinase under the α-myosin heavy chain (α-MyHC) promoter (Figure 1B).

**Figure 1:**
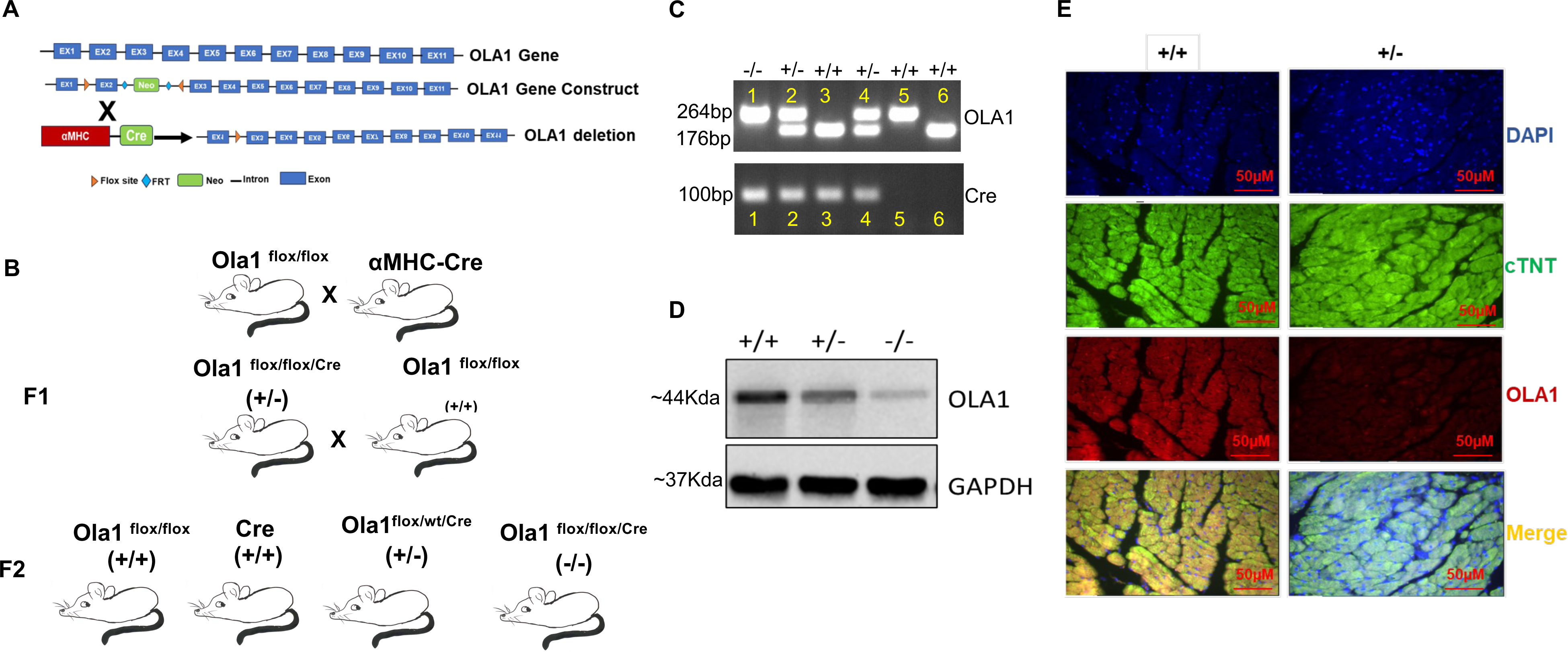
Generation of cardiac-specific OLA1 (cKO) knockout mice. A) The schematic diagram illustrates the OLA1 construct used to generate Ola1f/f (floxed) mice, which possess loxP sites flanking exon 2 of the Obg-like ATPase 1 (Ola1) gene. LoxP sites flank exon 2 of the OLA1 gene, and Frt sites flank the neomycin resistance (Neo) cassette. B) The schematic diagram shows the mice breeding strategy used to generate cardiomyocyte-specific OLA1 knockout mice. OLA1flox/flox mice were crossed with αMHC-Cre to allow cardiomyocyte-specific deletion of OLA1. C) The agarose gel electrophoresis image displays PCR amplification of approximately 176bp (Wild type), approximately 264bp (Flox), and 100bp (cre) genotypes. D) The Western blot shows the expression of OLA1 and GAPDH protein in +/+, +/- and -/- mice heart tissue, with GAPDH serving as a loading control. E) Immunofluorescence staining of heart sections from OLA1 haploinsufficiency (56 weeks) and control littermate (56 weeks) is shown, with Cardiac Troponin in Green, OLA1 in Red, DAPI in Blue, and Merge in Orange.

For genotyping, we used DNA from mouse ear punches taken at weaning. Amplified PCR products (using *Ola1* and Cre-specific primer sets) were visualized on 1.5% agarose gels to confirm their genotypes. Cardiac-specific homozygous (-/-, cKO) *Ola1* mutant mice showed a single band for flox-flox (264 bp) and Cre (100bp). Heterozygous (+/-, haploinsufficiency) mice samples displayed two bands (264bp for flox-flox and 176bp for wild type) along with the Cre band. Control (+/+) were considered as either flox or Cre positive littermates (Figure 1C). Interestingly, we observed that *Ola1* cKO homozygous pups (-/-) had developmental issues, resulting in fewer pups compared to heterozygous ones. We confirmed the activation of Cre recombinase, resulting in OLA1 deletion, by immunoblotting heart tissue lysates from homozygous (-/-) heterozygous (+/-, haploinsufficiency) and control (+/+) mice (Figure 1D). As expected, we found significant downregulation of OLA1 protein in the hearts of -/- and +/- mice as compared to +/+ controls. We further confirmed the deletion of *Ola1* in cardiomyocytes by staining heart sections with troponin (a cardiomyocyte marker) and OLA1. We found that the level of OLA1 expression was significantly down-regulated in troponin-positive cells, suggesting OLA1 deletion in cardiomyocytes (Figure 1E). These finding indicate that our model functions properly for further study.

### Cardiac-specific deletion of *Ola1* leads to increased mortality in mice

Previous studies on global Knockout mice have shown that homozygous OLA1 deletion is embryonically lethal, while heterozygous mice survive but develop severe phenotypes (5,11). To investigate its role in cardiac physiology, we generated *Ola1* cKO mice. Interestingly, we found that cardiac-specific *Ola1* homozygous (-/-) mice were also embryonically lethal, with only 4-5 homozygous (-/-) pups born, surviving up to 33 weeks (Figure 2A). Additionally, we that -/- mice had a smaller body size compared to +/+ mice (Figure 2B).

**Figure 2.**
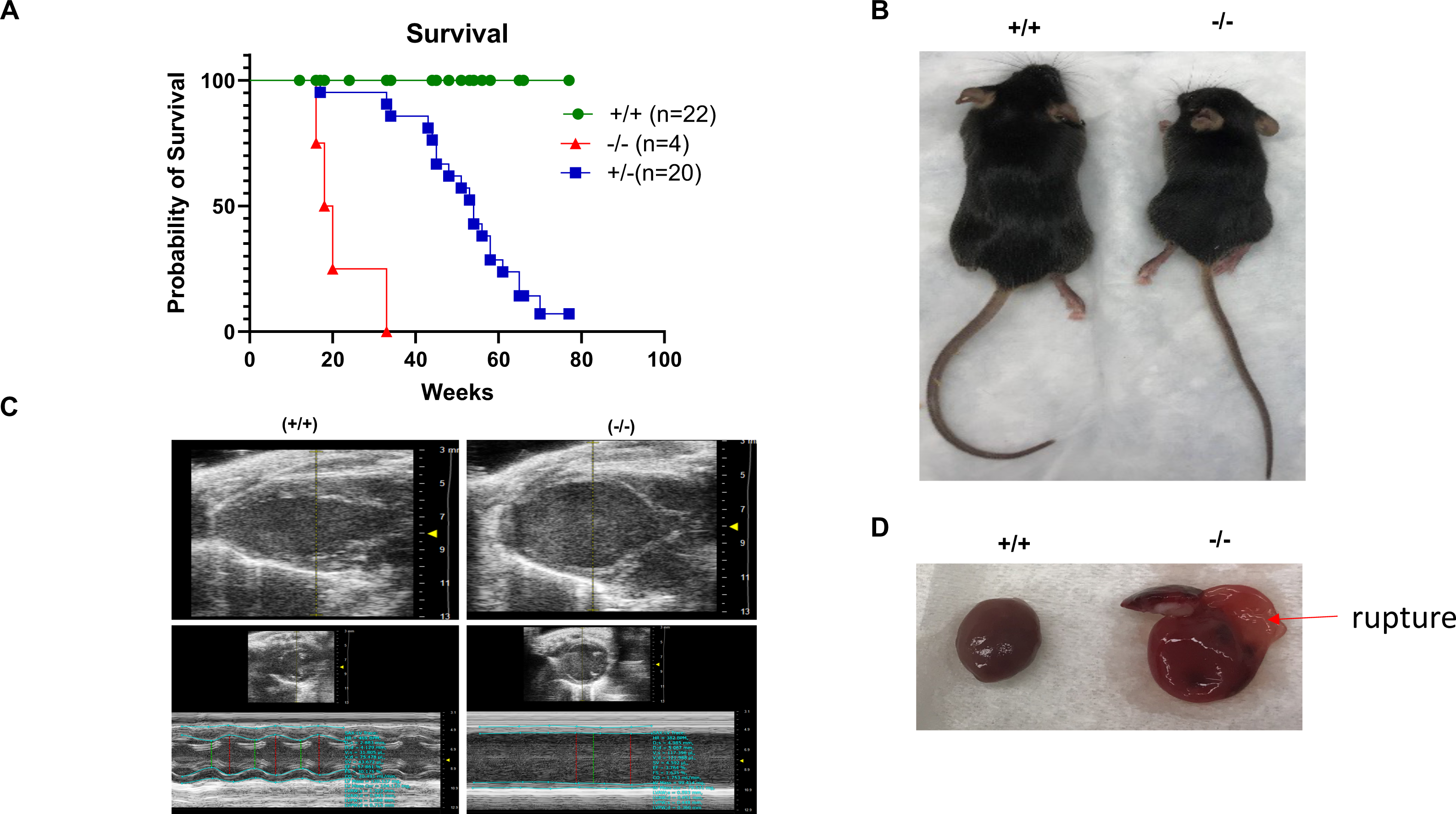
Survival study, Kaplan-Meier survival curves of OLA1-cKO and control littermate (Cre and Flox) mice. A) Mortality of homozygous OLA1-cKO (-/-, OLA1flox/flox X MHC-Cre) mice and OLA1 haploinsufficiency (+/- OLA1flox/WT X βMHC-Cre). A. OLA1 deficiency leads to death due to dilated cardiomyopathy in mice. B) Representative mice image of homozygous OLA1-cKO (-/-) and Control littermate (+/+) at the age of 5 weeks old. C) Representative images of mice echocardiogram showing cardiac dysfunction of OLA1 homozygous OLA1-cKO (-/-, age 14 weeks old), and control littermate OLA1 KO (+/+,14 weeks). D). Representative image of mice heart of OLA1-cKO (-/-, 14 weeks) and control littermate (+/+, 14 weeks). Arrow shows rupture of the heart.

To assess the impact of homozygous *Ola1* deletion on heart function and phenotype, we examined the mice at 14 weeks of age. We found that they developed severe cardiac dilation (Figure 2C), leading to cardiac rupture (Figure 2D). Given the importance of OLA1 in cardiomyocyte function, we monitored the survival of haploinsufficient (+/-) mice over an extended period. Mortality was significantly higher in +/- mice, with deaths starting at 33 weeks and peaking between 50-70 weeks of age (Figure 2A). These findings suggest that the *Ola1* gene plays a crucial role in cardiomyocyte physiology and the regulation of cardiac function.

### *Ola1* heterozygosity leads to dilated cardiomyopathy

Given our findings, we further studied *Ola1* haploinsufficiency mice. At a young age, we did not observe any discernible differences in phenotype or cardiac function between +/+ and +/- mice. However, as the mice aged, we observed that *Ola1* haploinsufficiency (+/-) mice spontaneously developed cardiac morphological changes and reduced body size in comparison to control littermates at same age (Figure 3A). We observed that OLA1 haploinsufficiency (+/-) mice developed cardiac dysfunction spontaneously, without any induced stress. These mice exhibited increased heart size and reduced body size in both male and female mice (Figure 3A-C). We quantified the heart weight/tibia length (HW/TL) ratio and body weight/tibia length (BW/TL) ratio in mice aged 50-75 weeks (Figure 3C-D).

**Figure 3.**
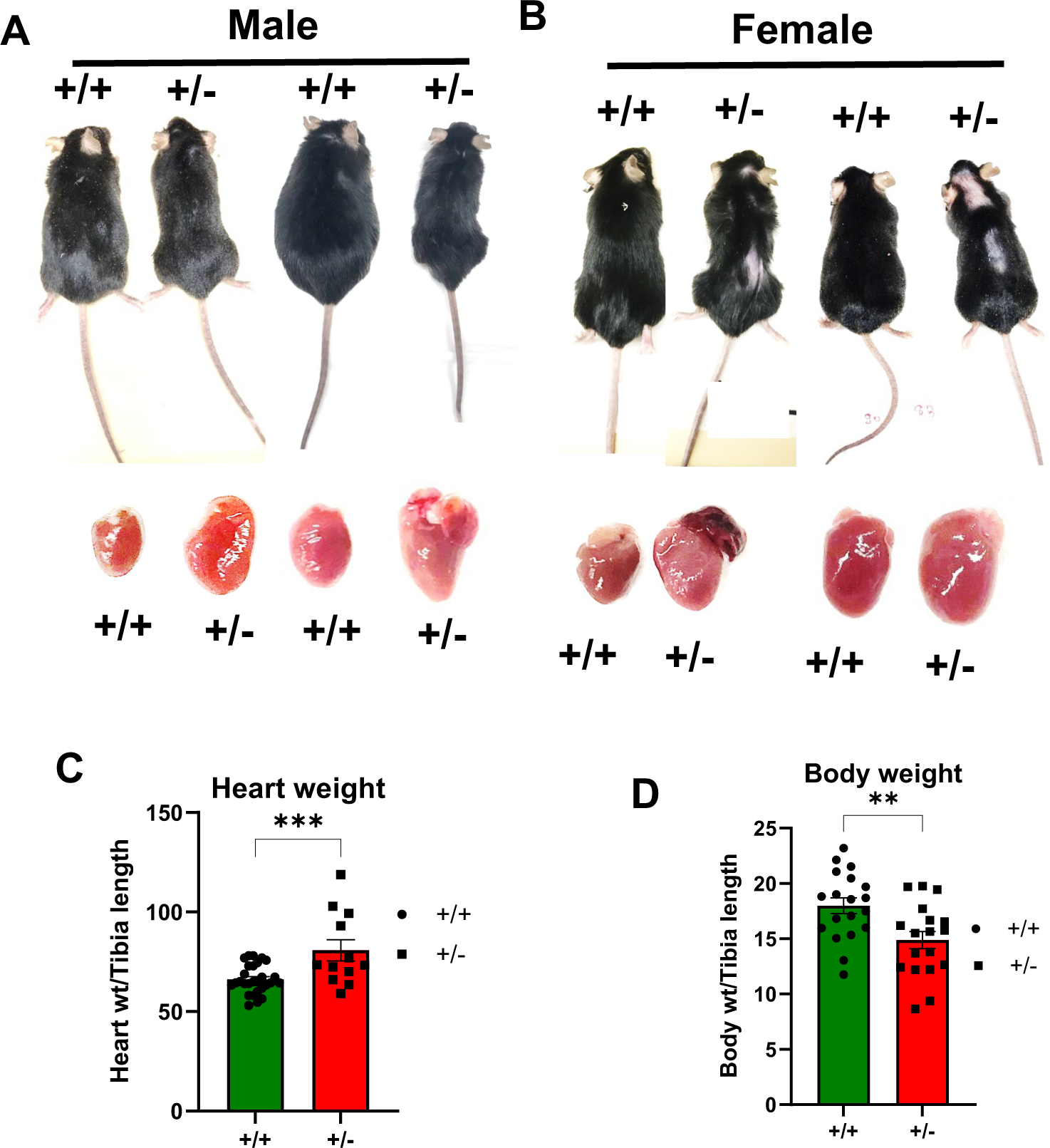
Cardiac function and morphology in aged males and females of OLA1 haploinsufficiency (+/-) and control mice (A, B) Representative images mice and isolated heart from OLA1 haploinsufficiency (+/- OLA1flox/WT X βMHC-Cre) and control littermate (+/+, OLA1flox/flox) age 50-65 weeks old. C,D) Quantifying the heart weight/tibia length (HW/TL) ratio and Body weight/tibia length (BW/TL) ratio in 50-75 weeks OLA cKO and control littermate mice. P>0.05 consider as significant.

### Assessment of cardiac function and structural remodeling in *Ola1* heterozygous mice

To understand whether OLA1 affects cardiac function and structural remodeling in vivo, we assessed LV function using echocardiography in both OLA1-cKO (+/-) and control littermate (+/+) mice. We allowed the mice to age naturally without cardiac stress until 80 weeks. Our findings indicate that the OLA1 haploinsufficiency mice (+/-) progressively developed severe ventricular dilation by 50 weeks, as evidenced by an increase in LV chamber diameter (Figure 4A). The dilation became more pronounced by 55 weeks, accompanied by a notable reduction in cardiac function, with a 10% decrease in ejection fraction (%EF) and fractional shortening observed in OLA1-cKO mice (Figure 4B-C). The LV chamber size was significantly larger and wall thickness was reduced in OLA1-cKO mice compared to control mice (Figure 4D-L). Also, the survival was significantly lower in haploinsufficiency (50% OLA1 knockout) mice compared to littermate control mice. Importantly, we observed rupture of the LV wall in the some of the deceased mice.

**Figure 4.**
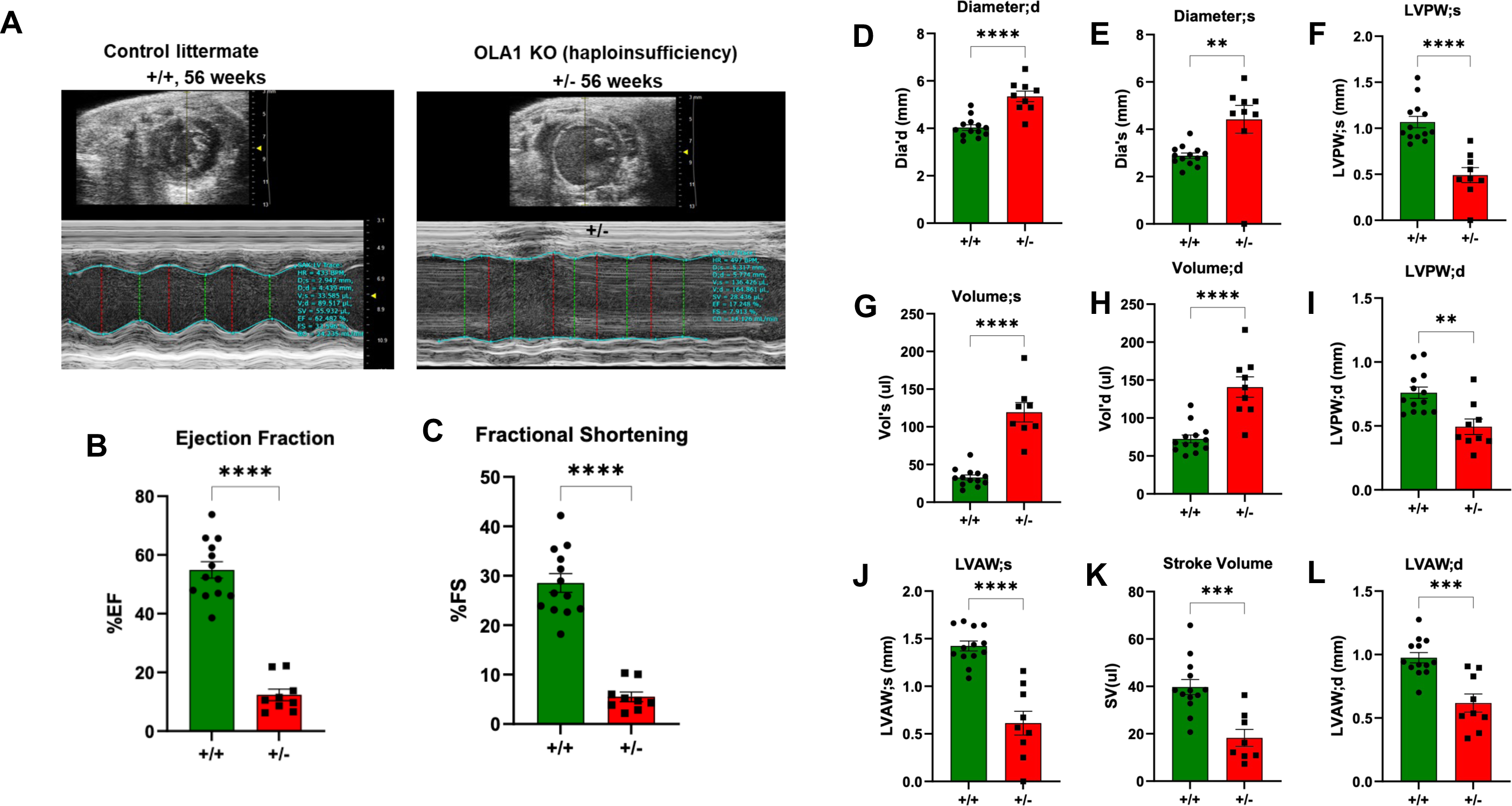
Cardiac function in aged OLA1 haploinsufficiency (+/-) and control mice. **(A)** Representative M-mode echocardiogram images showing cardiac dilation and dysfunction in OLA1 +/- mice (55 weeks). **B-L**) Comparative quantitative analysis of heart function among OLA cKO (+/-) and control littermate (+/+)., Ejection fraction (%EF), Fractional Shortening (%FS), Diastolic and systolic diameter (mm), left ventricular end-systolic diameter (LVED), posterior wall thickness in diastole (PWTd), interventricular septum thickness left ventricular in diastole (IVSTd), and fractional shortening (FS) in 50-65 Weeks-old OLA cKO mice and control littermate mice. P> 0.05 consider as significant.

### Histopathological changes in *Ola1*-cKO mice

Pathological examination of the hearts from *Ola1*-cKO mice revealed enlargement of the heart. Histology evaluation using H&E staining showed thinning of the LV wall and increased LV chamber size (dilatation) in *Ola1*-cKO mice (Figure 5A). Interestingly, Masson’s trichome staining revealed increased fibrotic tissue deposition in heart tissue sections of +/- mice compared to control littermates (+/+) (Figure 5B). These data suggest that *Ola1* gene deletion leads to the spontaneous development of dilated cardiomyopathy, adversely affecting cardiac function and survival.

**Figure 5:**
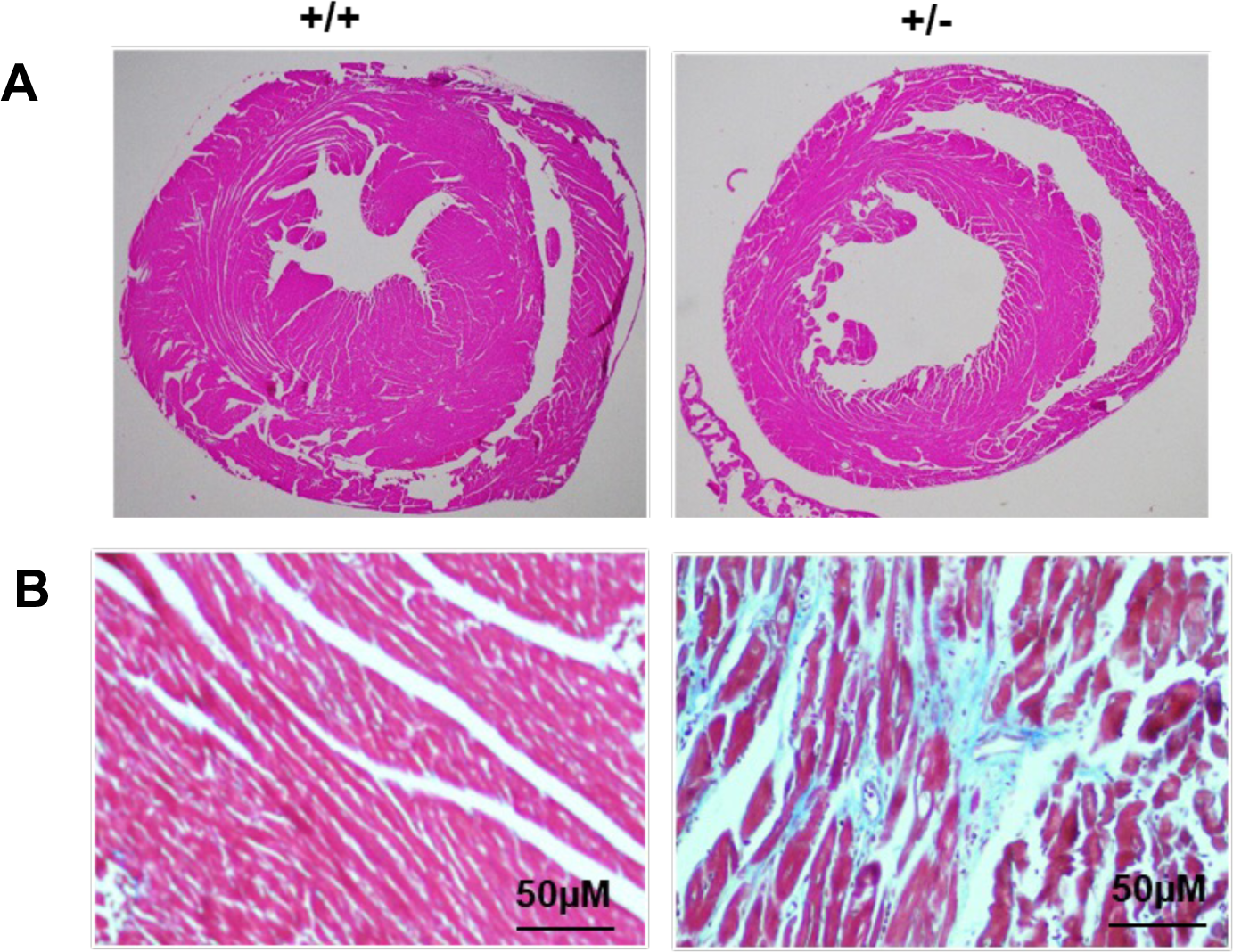
Hematoxylin and eosin (H&E) and Masson’s trichrome staining of mice hearts from OLA1 haploinsufficiency (55-65 weeks) and control littermates (55-65 weeks) (n=3) A) Hematoxylin and eosin (H&E) and b) Masson’s trichrome staining of heart sections isolated from OLA1 haploinsufficiency and control littermates.

### Loss of function for the Drosophila *ola1* gene affects pupal size and causes pupal lethality

The fruit fly, Drosophila melanogaster, is an excellent system for understanding gene-function in an organ-system. It is easy and inexpensive to maintain in a laboratory and offers tremendous advantages for genetic analyses. About 80% of known human disease genes have Drosophila homologs, and even those human disease genes that do not have a homolog can produce a phenotype like the human disease when expressed in flies (12). The Drosophila heart, also known as dorsal tube, is becoming a valuable model for studying heart development and genes that cause cardiomyopathies in humans (13–15).

We sought to examine the role of ola1 homolog in heart development and function using the fly system. The Drosophila heart starts to develop early during embryogenesis, which lasts for about 20 hours post fertilization. At the end of embryogenesis, fully developed embryos emerge or eclose as 1st instar larvae. After a day, they progress to the 2^nd^ instar, and after another day, they become 3^rd^ instar larvae, which lasts for 1.5 days (16). During these larval stages, they continuously grow, and the cells of the imaginal discs divide at a tremendous rate and number to produce adult structures in the pupal stage, which lasts for 4.5 days. Except for the digestive and nervous systems, which are retained into adult flies, all adult structures are generated from imaginal discs. During the larval stages, they feed continuously, but at the end of 3^rd^ instar larval stage, they stop feeding and attach themselves to a dry surface to pupariate.

The Drosophila *ola1* (CG1354) has about 74% amino acid identity with the human gene as well as with the mouse gene (human and mouse identity pegs at 99.5%). We utilized RNAi to determine the function of Ola1 in the heart by inducing the *ola1* RNAi with tinman (tinC-GAL4 driver; see Reference (17). This driver is only expressed in the heart, beginning with the cardioblasts in embryos (17). The induction of *ola1* with *tinC*-GAL4 driver resulted in pupal lethality, with 58% failing to emerge as adults versus 0% in the control. These dead or dying pupae, however, had a significantly reduced size, about 25% smaller than the control pupae (Figure 6A). This result shows that the size reduction in flies with the knockdown of *ola1* is consistent with the small size of mouse progeny that were homozygous for the loss of function of the *ola1* gene. There were escaper flies (42%) with normal-sized pupae, and the emerged adult flies did not show any measurable defects or phenotypes. These are likely cases where the *ola1* knockdown was not significant enough to cause a phenotype.

**Figure 6:**
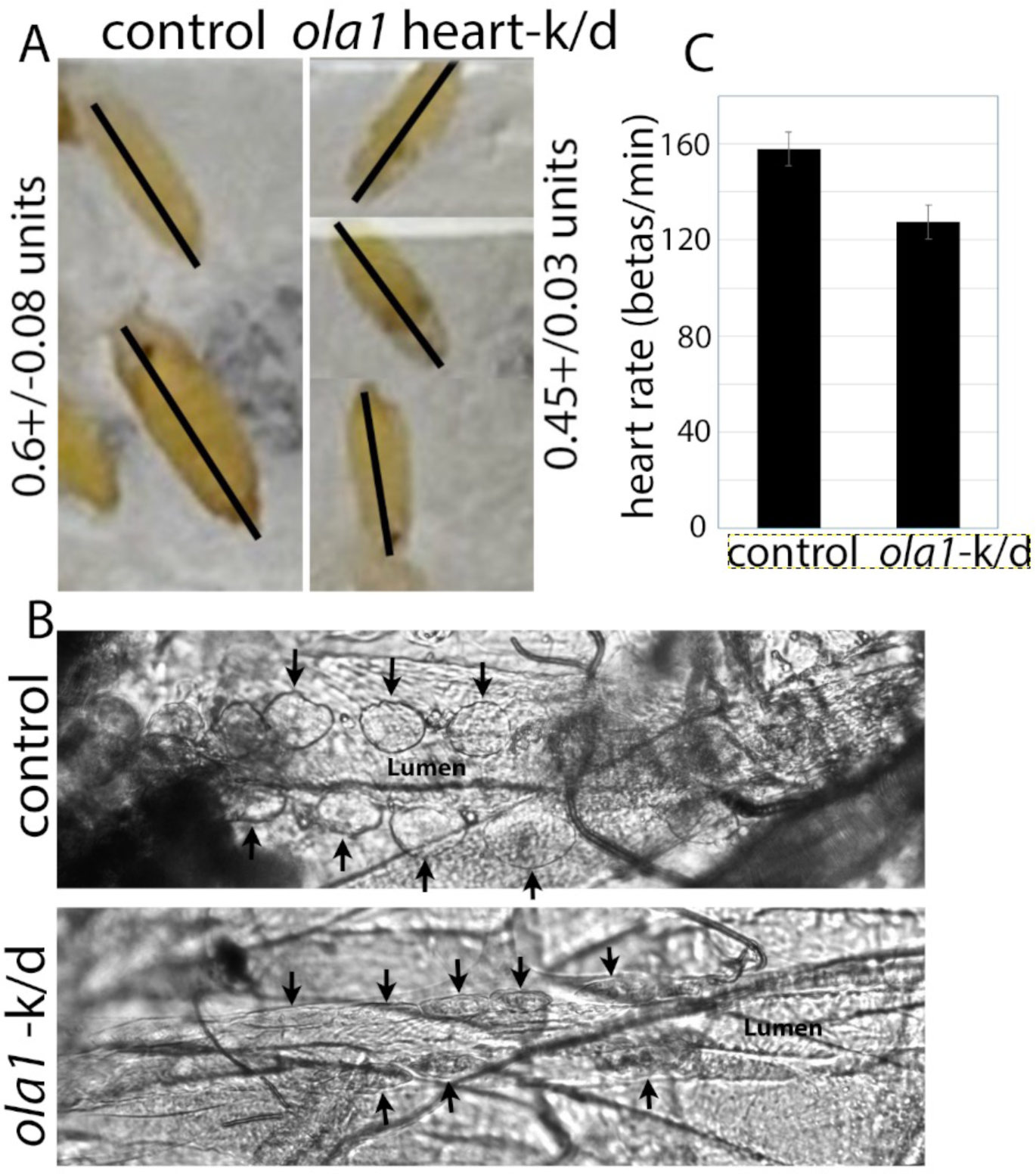
Heart-specific knockdown of *ola1*causes pupal lethality with heart defects in Drosophila. UAS-*ola1* RNAi transgene was induced with the heart-specific *tinC*-GAL4 driver. A) Normal pupae in control and the *ola1* RNAi, note the size difference between the two. B) Normarski images of heart from the control and the *ola1* RNAi larvae. Note the embedded round pericardial cells (arrows) in the control versus the elongated pericardial cells in RNAi larvae. C) The difference in the heart rate between control and the heart-specific RNAi of *ola1* larvae. The difference was statistically significant (see text for details). The images were obtained from live animals.

Next, we examined the heart in live 3^rd^ instar larvae from control and heart-specific *ola1* knockdown larvae (tinC>*ola1* RNAi). We monitored heart function by observing heartbeat for any aberrant rhythmicity under a microscope. In control larvae, the embedded pericardial cells lining the heart wall were well-defined and round, whereas in the *ola1* RNAi heart-specific knockdown larvae, the pericardial cells were elongated instead of round (Figure 6B). Consistent with this structural defect in the heart wall muscle cells, the heartbeats in *ola-*RNAi larvae appeared to be slow and abnormal (Movie *ola1-RNAi*) compared to the control (please see the attached movies files). We found that the larval heart beats in the control were 157+/- (N=10) per minute (N=10), whereas in the *ola1* heart-specific RNAi knockdown larvae, the heartbeats were 127+/-7 per minute (N=10). The difference was statistically significant student t test, P= 0.007) (Figure 6C). The beats were also asynchronous. These results confirm that *ola1* is essential for the proper development of the fly heart, including muscle structure, and for the proper heart function.

## Discussion

Heart failure remains a major health burden affecting more than 26 million individuals worldwide (1). Under physiological stress, the heart increases in size to reduce wall tension and maintain cardiac function. Sustained cardiac stress, however, leads to a pathological growth, progressing to dilated cardiomyopathy and heart failure(3). The major findings of the current study include: i) cardiac-specific *Ola1* knockout mice spontaneously developed severe left ventricular (LV) dilatation, thinning of the LV wall, reduced LV function, and, in some cases, ventricular wall rupture and death in mice, and ii) cardiac specific knockdown of Drosophila *ola1* causes pupal lethality with growth retardation, mis-shaped pericardial cells and asynchronous and slower heart rate in the larvae compared to the control.

*Ola1* has been shown to be involved in the translational regulation of cell proliferation, growth and protein synthesis in both mammals and yeast (5–7). Previous study from our lab has shown that upon ANG II administration in mouse, initially, OLA1 protein expression in the myocardium gradually increased at 3d and 14d (compensatory hypertrophic response), followed by a subsequent decline of OLA1 expression at 28 days (decompensatory phase) (8). These studies suggest that OLA1 is essential for adaptive response during the initial stages of heart disease, while a decrease in OLA1 could be a cause/effect of molecular switch from the initial protective compensatory hypertrophic response transitioning into maladaptive response.

Therefore, the objective of this study was to determine the consequence of cardiac-specific ablation of OLA1 on cardiac remodeling and function *in vivo*. OLA1 is essential for normal progression of mammalian development(18). In a previous report (5), OLA1 null mice showed growth retardation and developmental delay due to reduced rate of cell proliferation through translational regulation of p21 (5). In contrast, in the present study we observed that cardiac-specific OLA1 knockout mice are viable and exhibited no obvious morphological and functional abnormality before 6 weeks. However, as the mice aged, OLA1-cKO mice spontaneously developed cardiomyopathy. Cardiac-specific OLA1 gene ablation in mice resulted in the spontaneous development of severe LV dilatation, thinning of the LV wall, reduced LV function, and in some cases, ventricular wall rupture. In another on-going study in our laboratory, we have identified mutations in the OLA1 gene in heart tissue of patients with heart failure (9). These data suggest that OLA1 is critical for cardiac homeostasis and it’s deficiency adversely affects heart function.

The Drosophila heart is an excellent system to study the role of a gene in heart development and function (Selma-Soriano et al. 2021; Vogler and Bodmer, 2015; Reim et al. 2005). The fly *ola1* gene has about 74% protein homology to the vertebrate gene. Consistent with the conservation of gene function, knock down of *ola1* in developing heart cells resulted in pupal lethality with growth retardation (Figure 6A), which we attribute to the defects in pumping of nutrients. This reduced pumping was seen with a reduced and asynchronous heart rate (Figure 6C). The pericardial cells were significantly affected in terms of their structure, an uneven lumen (Figure 6B), and what appears to be dilated cardiomyopathy. Additional work is needed to determine if the cardiocytes were also affected. These results clearly show that *ola1* has a significant role in fly heart development and function and should contribute to our understanding of the function of *ola1* in heart across organisms.

It is unclear which signaling mechanisms are disrupted due to decreased OLA1 (as observed in heart tissue from heart failure patients) or loss of OLA1 (as in the cardiac-specific knockout mouse model). Previous studies have shown that OLA1 plays an important role in protein translation(6,18). Our previous study demonstrated that OLA1 modulated the in vitro response to ANGII treatment via the GSK3β/β-Catenin pathway(8). We anticipate that our on-going studies might identify the molecular mechanisms underlying the role of OLA1 in cardiac pathophysiology and DCM.

In conclusion, our study provides evidence that OLA1 plays a critical role in cardiac pathophysiology and that its deficiency leads to dilated cardiomyopathy and dysfunction. In our future studies, we will determine if targeting OLA1 rescues experimental dilated cardiomyopathy and what molecular signals are modified that could be beneficial in limiting the progression of heart failure.

## Supporting information

Supplement_Movie_control

Supplement_Movie_ola1RNAi

## Funding Support and Author Disclosures

This work was partly supported by US Department of Defense (DoD) CDMRP grant PR220330 (Dr. Krishnamurthy) and American Heart Association (AHA) Transformational Project Award (19TPA34850100, Dr. Krishnamurthy), National Institutes of Health grant R01NS131315 (Dr. Bhat). All other authors have reported that they have no relationships to disclose that is relevant to this study.

## Author contributions

PK and PKD conceptualized the initial project and study design. PKD performed the mouse experiments and KB lab members (HK and GKK) performed Drosophila experiments. PK and PKD prepared the initial manuscript draft. KB, HK, GKK and SS read and revised the manuscript draft and provided critical interpretations and insights. All authors discussed the results and interpretation and commented on the manuscript at all stages.

## Disclosures

None. All the authors have reported “nothing to disclose”.

---

**Supplementary Figure 1:**
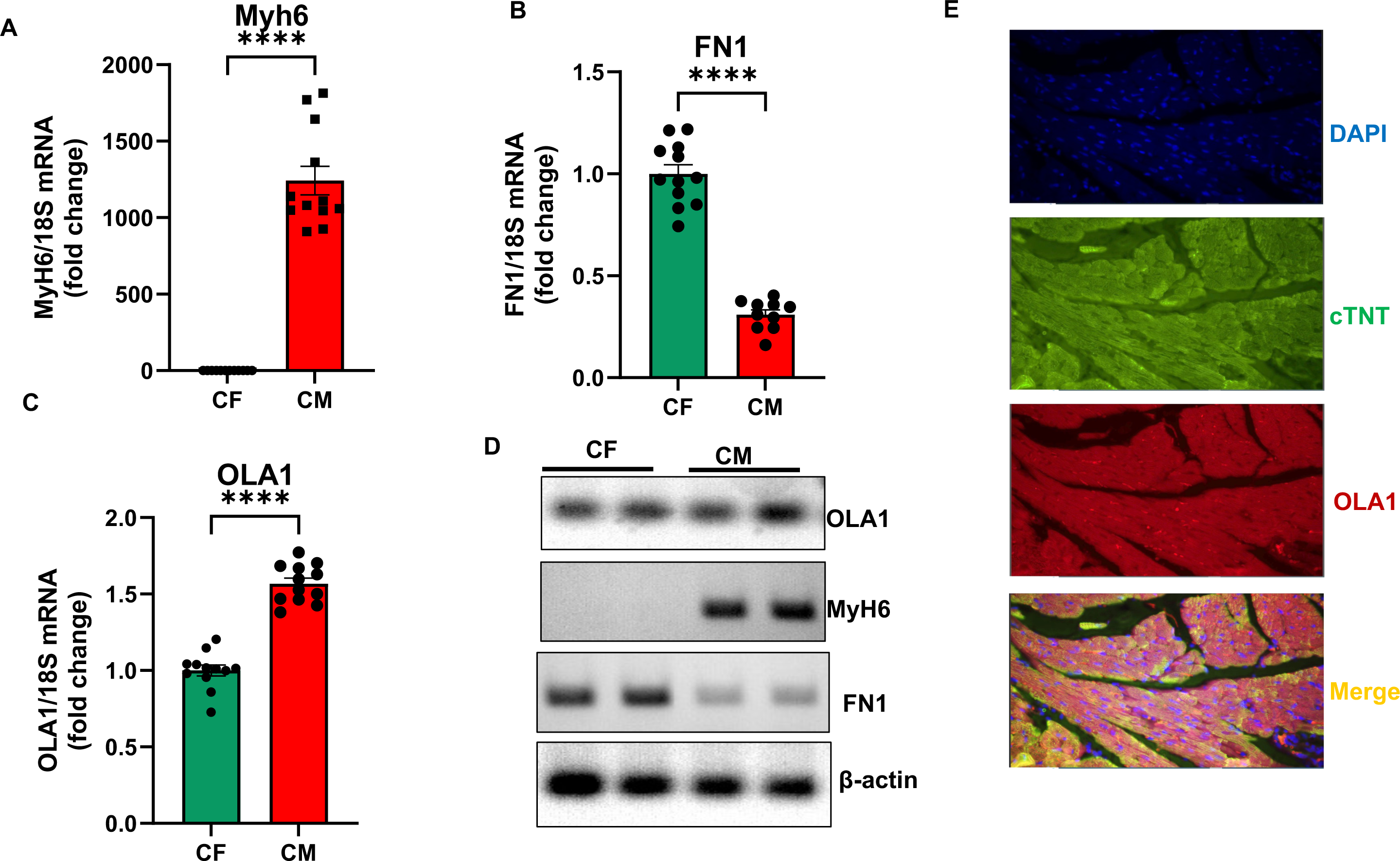
OLA1 expression in mouse heart tissue sections collected from young mice (12 weeks old). The qPCR assay measured mRNA expression of (A) Myh6 (Cardiomyocyte marker), (B) Fibronectin (Fibroblast marker), and (C) OLA1 expression in cardiac fibroblasts (CF) and cardiomyocytes (CM) isolated from adult mouse hearts. (D) Semiquantitative PCR. (E) Immunofluorescent staining of cardiac tissue showing OLA1 expression in cardiomyocytes. DAPI (Blue), Troponin T (Green), OLA1 (Red), Merge (Orange).

**Supplementary Figure 2:**
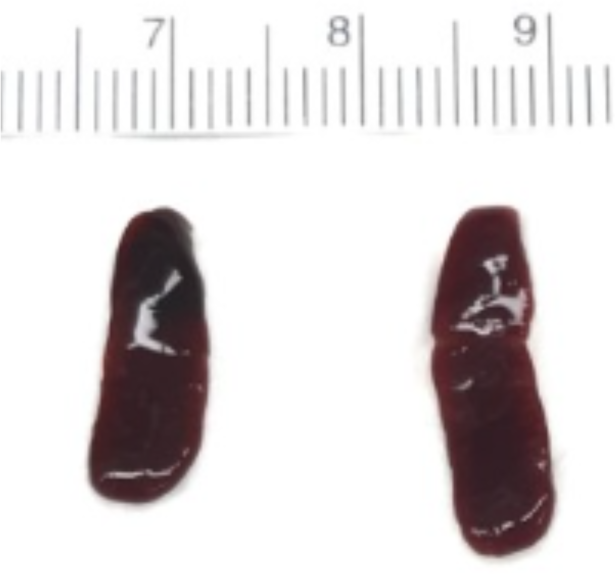
Representative image of a spleen isolated from OLA1 haploinsufficiency and control littermate weeks.

**Supplementary Movies**

**Control movie:** Larval heart was dissected, and the movie was obtained in live animals. Anterior end is to the right. Note the synchronous and rhythmic pattern of heartbeats.

***ola1* RNAi movie:** Anterior end is to the right. Note the slow heart-rate.

## Notes

### Competing Interest Statement

The authors have declared no competing interest.

